# clusterTools: proximity searches for functional elements to identify putative biosynthetic gene clusters

**DOI:** 10.1101/119214

**Authors:** Emmanuel LC de los Santos, Gregory L. Challis

## Abstract

**Motivation:** The low cost of DNA sequencing has accelerated research in natural product biosynthesis allowing us to rapidly link small molecules to the clusters that produce them. However, the large amount of data means that the number of putative biosynthetic gene clusters (BGCs) far exceeds our ability to experimentally characterize them. This necessitates the need for development of further tools to analyze putative BGCs to flag those of interest for further characterization.

**Results:** Clustertools implements a framework to aid in the characterization of putative BGCs. It does this by organizing genomic information on coding sequences in a way that enables directed, hypothesis-driven queries for functional elements in close physical proximity of each other. Genomic sequence databases can be constructed in clusterTools with an interface to the NCBI Genbank and Genomes databases, or from private sequence databases. clusterTools can be used either to identify interesting BGCs from a database of putative BGCs, or on databases of genomic sequences to identify and download regions of interest in the DNA for further processing and annotation in programs such as antiSMASH. We have used clusterTools to identify putative and known biosynthetic gene clusters involved in bacterial polyketide alkaoloid and tetronate biosynthesis.

**Availability and Implementation:** Clustertools is implemented in Python and is available via the AGPL. Stand-alone versions of clusterTools are available for Macintosh, Windows, and Linux upon registration (https://goo.gl/forms/QRKTkpqiA0g31IWp1). The source-code is available at https://www.github.com/emzodls/clusterArch.

**Supplementary information:** A manual describing the Python toolkit that powers clusterTools, as well as the HMMs constructed for the tetronate search is available online.

## 1 Introduction

Diminishing returns from screens of traditional chemical libraries coupled with the reduced cost of genome sequencing have fueled a renewed interest in mining genetic sequences for biosynthetic gene clusters (BGCs). This has led to the development of methods to analyze genome sequences to identify potential BGCs, most notably antiSMASH^1^. These have been successful in identifying BGCs, particularly those which contain polyketide synthase (PKS) or non-ribosomal peptide synthetase (NRPS) modules. However, the success of these algorithms in finding putative gene clusters has necessitated the need for further refinement and classification: the number of putative gene clusters exceeds our ability to experimentally characterize them. While advances in DNA synthesis, cloning, and heterologous expression will increase our ability to characterize a potential gene cluster, methods to select which clusters to prioritize are still needed.

One strategy in identifying BGCs of interest is to consider the functions of the different genes involved in a BGC. The natural products produced by a BGC contain specific chemical moieties that are synthesized from a limited set of bioavailable metabolites and identifiable enzymatic reactions. Taking advantage of this, and the fact that in bacteria, genes collectively involved in a specific function are usually in close physical proximity to one another, we can reduce the search space and identify regions of interest in the DNA. We do this by creating a “biosynthetic hypothesis” about how a natural product is produced, specifically the genes or functional domains involved in its biosynthesis. You can further narrow this space down by considering other functional elements such as regulatory or resistance genes. Querying genomic information in this targeted way, you can significantly reduce the portion of the data that requires further analysis, and increase the chances of finding putative BGCs of interest.

multiGeneBLAST^2^ is an open-source tool that can be used for this purpose. By reformatting FASTA headers of NCBI Genbank records, it identifies regions of DNA from databases that contain multiple BLAST^3^ hits near each other from a set of genes specified by the user. However, due to the repetitive nature and large size of PKS and NRPS genes involved in natural product biosynthesis, BLAST searches are oftentimes limited, producing many false positives. A solution to this is to instead consider the functional domains that a gene contains. These are identified through tools such as HMMer^3^ using profile Hidden Markov Models (HMMs). HMMer allows you to identify functional domains in a gene using custom-built HMMs for them. By searching for proteins in this way, we can identify uncharacterized genes that perform required functions of interest in your biosynthetic hypothesis in addition to known, well-characterized proteins. By combining BLAST and HMMer searches, one has more flexibility and can increase the granularity of their searches.

Here we present clusterTools, software developed for this purpose. clusterTools establishes a framework to store information about a gene in a way that it is amenable to hypothesis driven queries using BLAST or specific domain compositions in the form of HMM Rules. This allows us to find sets of genes in close physical proximity that match a biosynthetic hypothesis. clusterTools is designed to work with existing genome mining software, allowing you to create sequence databases from antiSMASH output, or from NCBI Genbank and Genome queries. Databases can also be constructed from annotated genbank sequences in local databases. The interface to the NCBI allows you to download sequence information for hits to your queries that are in the NCBI. These putative gene clusters can then be further analyzed with other tools.

## 2 Implementation

### 2.1 Python Toolkit

The core of clusterTools is a Python toolkit that consolidates information from different sources to create Protein and Cluster python objects. Protein objects will contain information concerning the physical location on the DNA of the sequence, and results of BLAST and HMMer queries of the protein. Cluster objects group Proteins in functional units. A set of functions was written to manipulate and compare these objects.

Notable functions include the parsing of HMMer outputs to annotate the functional domains in a protein based on HMM hits. The criteria for annotating the region of the protein with a specific function are based on its HMMer score (where a higher score takes precedence), coverage, and minimum domain size. Results from different HMM searches can be merged in this way following the same criteria. When two HMMer hits of the same type are beside each other, these are merged when they meet specific coverage criterion. Annotating proteins in this way allows you to extract domain strings and sets of Clusters and Proteins, which are representatives of the functional elements contained in a protein or a cluster and the order in which they occur.

An all-v-all BLAST can be performed of proteins and clusters stored in this toolkit. This enables you to calculate metrics such as distances and between a protein and a cluster based on their BLAST bitscore. These distances can be used for visual comparison in comparing clusters of interest.

While the GUI provides access to some of these functions, the Python toolkit can also be used separately in scripts for even more specific cluster analyses and comparison. A full description of the functions included in the toolkit and their use can be found in the SI.

### 2.2 Graphical User Interface

A GUI is implemented to provide users access to some of the functions of the Python toolkit. The GUI has three parts: (1) Prepare Database, (2) Run Search, and (3) Load Previous Search.

#### 2.2.1 Prepare Database

This provides functionality for the creation of sequence databases that can be processed by clusterTools. clusterTools databases are protein sequences in FASTA format whose headers are specifically formatted with information about the protein identity, source, and physical location on the DNA. Databases can be constructed from annotated genbank files that are downloaded from the NCBI, or from private sequencing projects. An option also exists to create a custom database from a subset of the NCBI Genbank or Genomes databases. Databases created from antiSMASH cluster genbank files will contain information about their predicted product as assigned by antiSMASH. This functionality allows you to quickly identify antiSMASH annotated clusters that contain functionality that you are looking for.

#### 2.2.2 Run Search

Once a database has been constructed, it can be queried for functional elements based biosynthetic hypothesis, regulatory, or resistance genes. clusterTools accepts protein FASTA files as input sequences for genes. Alternatively, a protein coding sequence can also be inputted manually and be used in a query. clusterTools will accept hmms constructed with hmmbuild^4^ (<= v3.1 in Linux/OS X, <= 3.0 in Windows) as input. An interface to hmmbuild is provided in the Linux and OS X versions, allowing the construction of custom HMMs using a single protein sequence or a multi-sequence alignment in FASTA or Stockholm format

The user can then query the database for regions of DNA that contain BLAST hits of specified genes, or genes that satisfy HMM logic rules built from loaded HMMs, creating a search list of functional elements. Items in the search list can be required or optional. To register as a hit, a region of DNA needs to contain all the required hits and some optional hits so that the total number of hits in the region is greater than or equal to the minimum number of hits specified. This is useful in the case where you have alternative biosynthetic hypotheses, when there is uncertainty about what kind of gene is involved in a particular step of the pathway, or if certain genes are implicated in the biosynthesis of a natural product of interest but are not required. clusterTools will report any hit that is in the search list regardless of whether or not they are required. The window size of the DNA the hits need to be in, the minimum domain size, and the minimum E-Values for the homology searches can be changed adjusted to fine tune the query.

Clustertools will perform the requested search, running BLAST on the database with the specified proteins as queries, and hmmscan^4^ with the chosen HMMs. Output from these searches are used as input to functions in the toolkit to identify regions of DNA that correspond to the search. clusterTools then summarizes the results providing the user with the regions of DNA that contain elements of the search list, as well as the coding sequences that correspond to each of the hits. A webpage visualizing the results of a clusterTools search is also generated. If the database queried contains sequences from the NCBI, the user has the option to download the region of DNA for further analysis. Figure 1 summarizes a proposed clusterTools workflow. The user has the option to save the results of a search for future use.

**Figure 1.**
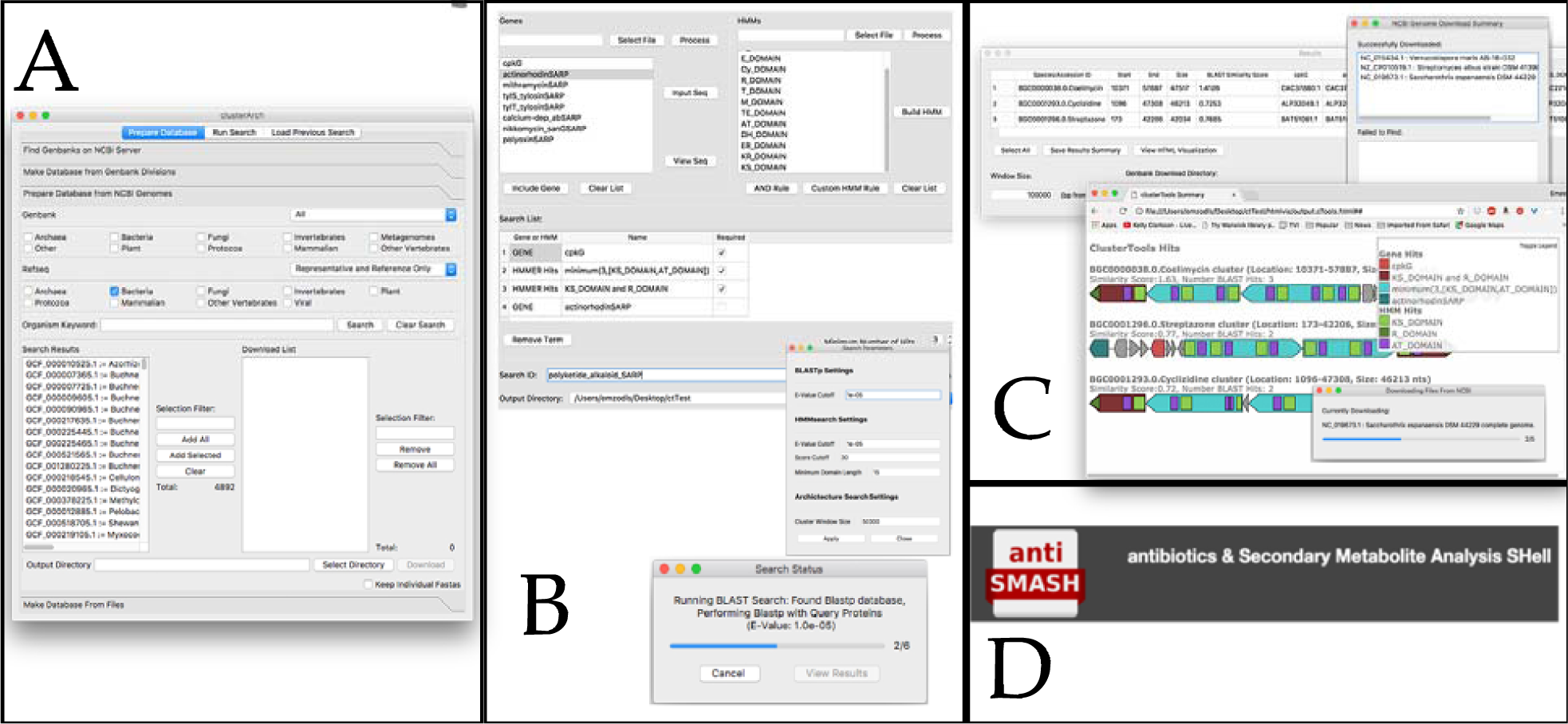
clusterTools workflow. A typical clusterTools workflow involves. (A) the creation of a database to conduct searches, (B) specification of search conditions with that include a combination of genes and functional domains, (C) Analysis of the results and selection of hits for download, (D) further analysis of sequences of interest.

#### 2.2.3 Load Previous Search

By saving the results and search information from a previous search, a user can quickly conduct a related search, changing the search list by altering the required genes or specifying a different set of HMM Rules from the HMMs used in the initial search. Certain search parameters such as the window size can also be changed. Since the BLAST and hmmscan steps are skipped and previous results are used, searches done this way are completed much more rapidly. Functionality to download sequences as a result of these searches is maintained in this mode.

## 3 Results

### 3.1 Polyketide Alkaloids

With our studies on the biosynthesis of coelimycin P1 and reports on the biosynthesis of other polyketide alkaloids, we hypothesized that polyketide alkaloid biosynthesis in actinobacteria go through universal steps; specifically, reductive chain release and transamination^5^. We queried Genbank and Refseq databases with the following search list:

**Table 1.** Polyketide Alkaloid Search.

| Gene or HMM Name |  | Required |
| --- | --- | --- |
| Gene | cpkG (Accession: NP_630377) | Yes |
| HMM | (KS Domain and AT Domain) | Yes |
| HMM | (KS Domain and R Domain) | Yes |

**Figure 2.**
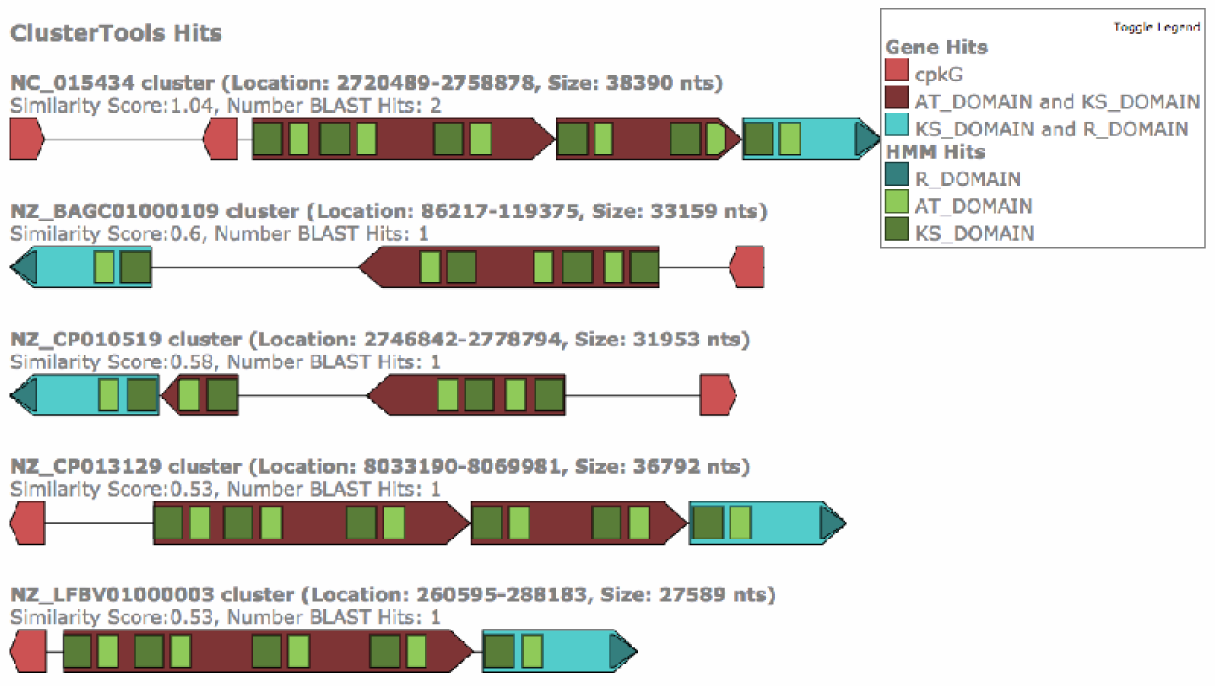
Visualization of Polyketide Alkaloid Search. HTML output showing some of the results of the polyketide alkaloid search. The query was conducted on the Refseq representative bacterial genomes from the NCBI.

This search finds stretches of DNA that contain at least two polyketide modules one of which containing a reductive chain release domain, near a transaminase gene. We used the cpkG gene sequence from *S.coelicolor* to search for putative □-transaminases and NRPS/PKS HMMs from Bachmann *et al^6^.* for the functional domain search. Our search resulted in 21 putative BGCs whose sequences we downloaded and further analyzed with antiSMASH and manual annotation. We predicted that some of these clusters will produce novel polyketide alkaloids, while some of them are predicted to produce known natural products, or derivates of these. The results of this search and subsequent analyses are described in Awodi *et al^4^.*

### 3.2 Tetronates

Extensive work has been done on the biosynthesis of tetronates, this is summarized in Vieweg *et al*^7^. Briefly, biosynthesis of the tetronate requires three genes: (1) FkbH-type gene that catalyzes the loading of 1,3-bisphosphoglycerate (1,3-BPG) into an acyl-carrier protein (ACP), (2) ACP gene that 1,3- BPG is loaded into, (3) standalone FabH-type gene that catalyzes the condensation of 1,3-BPG to a polyketide chain. Other genes that are associated with tetronate biosynthesis are: (1) an acetylase that acetylates a hydroxyl-group in the tetronate, (2) a dehydratase that eliminates the acetyl group forming an exocyclic double bond, and (3) a diels-alderase, that performs a diels-alderase reaction resulting in a spirotetronate.

Using the sequences of genes from known tetronates and spirotetronates, we constructed HMMs to search for putative BGCs that produce tetronates. In this case we used the following search conditions:

**Table 2.** Tetronate Search.

| Gene or HMM | Name | Required |
| --- | --- | --- |
| HMM | ACP | Yes |
| HMM | FabH | Yes |
| HMM | FkbH | Yes |
| HMM | acetylase | No |
| HMM | dehydratase | No |
| HMM | Diels-alderase | No |

In order to validate the search conditions, we first tested it on a clusterTools database constructed from the MiBIG^8^ database, which is a database that contains known biosynthetic gene clusters. Table 3 shows the results of the MiBIG query this corresponds to the known tetronate BGCs that are in MiBIG. Having validated the HMMs we conducted a similar search on a database containing reference and representative bacterial genomes from the NCBI genomes database. The search yielded 48 putative tetronate BGCs. The HMMs used in this search and the hits from the NCBI Genomes search are included in the SI.

**Table 3.** MiBIG Tetronate Search Hits.

| Species/Accession ID | Start | End | Size | fabH | fkbH | acp | acetylases | dehydratase | diels_ald_all |
| --- | --- | --- | --- | --- | --- | --- | --- | --- | --- |
| BGC0000140.0.RK-682 | 9192 | 12387 | 3196 | ACZ65477.1 | ACZ65478.1 | ACZ65479.1 |  |  |  |
| BGC0000162.0.Tetrocarcin | 77303 | 82324 | 5022 | ACB37745.1 | ACB37746.1 | ACB37747.1 | ACB37748.1 | ACB37748.1 |  |
| BGC0000082.0.Kijanimicin | 32122 | 37107 | 4986 | ACB46483.1 | ACB46482.1 | ACB46481.1 | ACB46480.1 | ACB46480.1 |  |
| BGC0000001.0.Abyssomicin | 10118 | 15057 | 4940 | AEK75497.1 | AEK75498.1 | AEK75499.1 | AEK75500.1 | AEK75501.1 |  |
| BGC0001183.0.Lobophorin | 74315 | 79269 | 4955 | AGC09489.1 | AGC09490.1 | AGC09491.1 | AGC09492.1 | AGC09492.1 |  |
| BGC0001004.0.Lobophorin | 28553 | 33514 | 4962 | AGI99492.1 | AGI99491.1 | AGI99490.1 | AGI99489.1 | AGI99489.1 |  |
| BGC0000036.0.Chlorothricin | 99866 | 105801 | 5936 | AAZ77702.1 | AAZ77703.1 | AAZ77704.1 | AAZ77705.1 | AAZ77706.1 | AAZ77701.1 |
| BGC0000133.0.Quartromicin | 22952 | 70230 | 47279 | AFI57023.1 | AFI57024.1 | AFI57025.1 | AFI57026.1 | AFI57004.1 | AFI57012.1 |
| BGC0001204.0.Versipelostatin | 60902 | 95798 | 34897 | BAQ21951.1 | BAQ21955.1 | BAQ21954.1 | BAQ21953.1 | BAQ21952.1 | BAQ21945.1 |
| BGC0001288.0.Maklamicin | 59328 | 68189 | 8862 | BAQ25501.1 | BAQ25502.1 | BAQ25503.1 | BAQ25504.1 | BAQ25505.1 | BAQ25497.1 |

## 4 Conclusion

We routinely use clusterTools to perform specific hypothesis-driven searches to identify putative BGCs of interest. While searches usually involve genes involved in the biosynthesis of a chemical moiety of interest, we have also reduced the search space by including elements of the cluster not involved in the biosynthesis, such as pathway specific activators or resistance genes. We believe that clusterTools complements existing software in increasing the power of *in silico* screening of BGCs of interest and reducing the number of potential BGCs that require further characterization to a more tractable number.

## References

1. Weber T, et al. (2010). antiSMASH 3.0 - a comprehensive resource for the genome mining of biosynthetic gene clusters. (2015) Nucleic Acids Res. 43: W237–W243.

2. Medema MH. (2012) Multigeneblast: Combined Blast Search For Multigene Modules. http://multigeneblast.sourceforge.net. N.p.. Web. 21 Mar. 2017.

3. Camacho C, Coulouris G, Avagyan V, Ma N, Papadopoulos J, Bealer K, & Madden TL (2009). BLAST+: architecture and applications. BMC Bioinformatics, 10, 421. http://doi.org/10.1186/1471-2105-10-421

4. Eddy SR (2011) Accelerated Profile HMM Searches. PLOS Computational Biology 7(10): e1002195. doi:10.1371/journal.pcbi.1002195

5. Awodi U, et al. (2017). Thioester reduction and aldehyde transamination are universal steps in actinobacterial polyketide alkaloid biosynthesis. ChemicalScience. 8. 411–415

6. Bachmann BO and Ravel J. (2009). In SIlico Prediction of Microbial Secondary Metabolic Pathways from DNA Sequence Data. Methods in Enzymology. 458:181–217.

7. Vieweg L, et al. (2014). Recent advances in the field of bioactive tetronates. Natural Product Reports. 31:1554–1584

8. Medema MH, et al. (2015). Minimum information about a biosynthetic gene cluster. Nature Chemical Biology. 11:625–631

